# Influence of co-occurring weakly pathogenic bacterial species on bacterial spot disease dynamics on tomato

**DOI:** 10.1101/2023.04.25.538297

**Authors:** Shreya Sadhukhan, Marie-Agnes Jacques, Neha Potnis

## Abstract

Mixed infections caused by multiple pathogenic and/ weakly pathogenic strains inhabiting the same host plants are common in nature and may modify pathogen dynamics. However, traditional plant pathogen studies have mostly focused on the binary interaction between a single host and a single pathogen. In this study, we have looked beyond this binary interaction and evaluated the impact of co-infection on disease dynamics on tomato using bacterial spot pathogen, *Xanthomonas perforans* (*Xp*) and two co-occurring weakly pathogenic strains of *X. arboricola* (*Xa*) and *Pseudomonas capsici* (*Pc*). Time-series greenhouse co-infection experiments monitoring disease severity and within-host population dynamics revealed higher disease severity in co-infection by three species compared to infection by *Xp* alone. However, co-infection by dual species, *Xp* and *Pc* or *Xa* resulted in lower disease severity compared to *Xp* alone. Thus, co-infection outcomes depend on interacting species. Weak pathogens could exploit *Xp* to colonize the host plant as indicated by their higher populations in co-infection. However, *Xp* population dynamics was dependent on co-infecting partner. While resource competition might be a possible explanation for lower *Xp* population in dual co-infection, interaction of *Pc* with the host was found to influence *Xp* population. Interestingly, *Xp* population was higher in presence of three-species interaction compared to *Xp* and *Xa* co-infection, suggesting potential modulation of co-operative interactions among *Xp* and *Xa* in three-species co-infection rather than competitive interactions. Humidity played a significant role in population dynamics of the three species. Overall, this study highlighted importance of co-infection dynamics in studying plant disease outbreaks.

## Introduction

Co-infections by multiple pathogenic species or genera are common in agricultural fields (Susi et al. 2015, 2022; Lamichhane and Venturi 2015; Abdullah et al. 2017). However, molecular, epidemiological and disease management studies have largely focused on binary interactions among dominant pathogen and the plant, with the design of mitigation practices targeting the dominant pathogen. Such limited approach not only ignores the importance of co-occurring pathogens or opportunistic pathogens that can likely contribute towards disease epidemiology but also rarely recognizes the ecological interactions among the dominant pathogen and the co-occurring weak pathogens co-occurring taxa that may alter dominant pathogen population dynamics (Baldotto and Olivares 2008; Sturz et al. 2000). With the growing appreciation of the importance of microbiota in extending plant immunity towards pathogen and in mediating interactions with the pathogen, recent studies are now focusing on how dominant pathogen and microbiota interactions may influence the overall disease outcome and pathogen population using approaches of amplicon sequencing or metagenome analysis (Wang et al. 2023; Newberry et al. 2020). Although these studies look beyond binary interactions of dominant pathogen and the plant, they still ignore the ecological dynamics of weak pathogens, rely on co-occurrence based on relative abundance values to derive interaction networks, and fail to tease apart the nature of true interactions that can influence dominant pathogen population and overall disease outcome.

It is largely assumed that mixed infections with multiple microbial species lead to the development of severe disease symptoms (Lamichhane and Venturi 2015) compared to single infections. However, this might not always be true (Stubbendieck and Straight 2016). The influence of co-infections on overall plant susceptibility may depend on various factors. First and foremost, of which, are microbe-microbe interactions. Interactions of the pathogen with the resident microbes including those with co-occurring opportunistic pathogens may range from antagonism, mutualism, cooperation, or synergism (Abdullah et al., 2017, Lamichhane & Venturi, 2015). These interactions may be mediated by resource availability, niche occupation or via production of compounds involved in synergistic or antagonistic interactions (Cornforth and Foster 2013). In a mixed infection, bacterial populations can co-operate by sharing virulence factors, siderophores, toxins, exo-enzymes, and bio-surfactants. For example, iron is a limited resource in the host environment. Competition for iron acquisition is considered as a reason for bacterial species to compete in the host environment. Interaction between different bacterial species influences their lifestyle and how they communicate. Inter-species quorum sensing is a possible mode for communication between different bacterial species who are in proximity (Rezzoagli et al. 2020). Opportunistic pathogens and other co-occurring microbes might exploit these characteristics in mixed infections. In case of plant-microbe interactions, plant defense response plays an important role, that adds a layer of complexity to the microbe-microbe-plant interactions. Plants can recognize pathogens by two strategies: microbial elicitors like pathogen-associated molecular patterns (PAMPs) or damage-associated molecular patterns (DAMPs). These patterns are recognized by plant receptors proteins called pattern recognition receptors (PRRs). Stimulation of PRR induces PAMP-triggered immunity (PTI) in host plants (Dodds and Rathjen 2010). The second method of detection is recognition by intracellular receptors of pathogen virulence molecules known as effectors, which induces effector-triggered immunity (ETI) in the host (Dodds and Rathjen 2010). In case of co-infection or mixed infection, the plant host’s defense mechanism might prioritize to invest its defense metabolites against certain pathogens. The prioritization depends on the mode of action of the pathogen (Castrillo et al. 2017; Hacquard et al. 2016). This raises the possibility that infection by one pathogen can impact the plant’s defense for the subsequent infection by a different pathogen or not (Abdullah et al. 2017). Alternatively, interaction of one of the coinfecting member with the host, such as induction of immune response, may alter the colonization by the other coinfecting member. This type of competition may exist between co-infecting pathogen species is referred to as apparent competition (Kinnula et al. 2017; Read 2001).

The disease outcome of co-infection depends on the extent to which each of the pathogen colonizes the host. It also depends on the timing of colonization (Benítez et al. 2013). Single inoculation time points have been used for most of the co-infection studies. Therefore, the possible influence of timing has not been clearly understood. In natural or field conditions sequential co-infections are more likely to be present rather than simultaneous co-infection. Studies reveal that sequential and simultaneous co-infections led to differences in disease severity (Marchetto and Power 2017).

Bacterial spot disease caused by *X. perforans* (also referred to as *X. euvesicatoria* species complex) is endemic to the southeastern US. Our previous metagenomic surveys indicated that co-occurrence of *Pseudomonas* spp. and *Xanthomonas* spp. is common in the bacterial spot infected tomato phyllosphere in Alabama fields (Newberry et al. 2020). Pectolytic, opportunistic, avirulent *Xanthomonas* species have also been reported to be isolated from mixed infections with *Pseudomonas syringae* from pepper and tomato transplants (Gitaitis et al. 1987). Similar reports of association of *Xanthomonas arboricola* with bacterial spot infected tomato and pepper have been noted in the subsequent years (Mbega et al., 2012, Myung et al., 2010, Newberry et al. 2020). Surveys carried out in soil-grown tomatoes in Sicily (Italy) showed that *Pseudomonas* spp. strains were frequently associated with *Xanthomonas perforans*, causal agent of tomato pith necrosis. These *Pseudomonas* spp. significantly increased pith necrosis and vascular discoloration symptoms when co-inoculated with *X. perforans* on tomato plants. In these co-inoculations, *X. perforans* population density was significantly higher than that of *X. perforans* inoculated individually (Aiello et al. 2017). Among the co-occurring members identified with *X. perforans* and those shared among tomato, pepper and solanaceous weed species included *X. arboricola* and *P. cichorii*-like species (Newberry et al. 2020). Indeed, report of *P. cichorii*-like strains, belonging to a novel species *P. capsici*, (Zhao et al. 2021) has suggested these pseudomonads may act as weak pathogenic species. *X. arboricola* has been referred to as opportunistic pathogen as outbreaks caused by them have been sporadic and dependent on conducive environmental conditions (Gitaitis 1987; Roach et al. 2018, 2019). The variation in bacterial spot disease severity or aggressiveness in the field conditions is often thought to arise from the changes in the dominant pathogen population structure or correlated with presence of certain pathogen genotypes carrying specific virulence factors. However, the extent to which co-infecting pathogens or weak pathogens may contribute towards disease outbreaks and explain severe outbreaks in the field conditions is still unclear (Potnis 2021).

In this study, we addressed this question by investigating the influence of co-infection by *X. perforans*, and other co-occurring weak pathogens, *X. arboricola* and *P. capsici* on overall disease severity on tomato and on population dynamics of a dominant pathogen, *X. perforans* and other co-occurring weak pathogens. We hypothesize that co-infection leads to overall higher disease severity and alters dominant pathogen population dynamics. Since high humidity conditions are known to favor colonization by weak pathogens, we also evaluated the influence of humidity on co-infection dynamics, both in terms of disease severity as well as population dynamics. Finally, we speculate on possible mechanisms to describe overall findings of higher disease severity but lower dominant pathogen population under co-infection.

## Materials and Methods

### Plant material and growth conditions

Tomato plants of cultivar FL47R, grown in greenhouse, were used for this study. After the seeds germinated, two weeks old seedlings were transplanted into 4” plastic pots with potting mix. Plants were kept in greenhouse at 28°-30°C under greenhouse conditions for 4-5 weeks before using them for our study.

### Bacterial strains, media, and growth conditions

Bacterial strains and plasmids used in this study are listed in Table 1. Bacteria were routinely cultured on nutrient agar (NA, Difco) supplemented with 15 μg/ml gentamicin or tetracycline or chloramphenicol and 50 μg/ml streptomycin where appropriate. These strains were grown for 24-48 hours at 28°C. The wild type bacterial strains *Xanthomonas perforans* AL65 (referred to as *Xp* hereafter), *Xanthomonas arboricola* CFBP 6826 (referred to as *Xa* hereafter), *Pseudomonas capsici* 93B.260 (referred to as *Pc* hereafter), were tagged with plasmids carrying distinct antibiotic resistance marker (Schlechter and Remus-Emsermann 2019).

**Table 1:** Bacterial strains and plasmids used in this study

| Strain | Short name; Features | Source |
| --- | --- | --- |
| *Xanthomonas perforans* AL65 | Xp AL65 WT, Streptomycin resistant | Infected pepper leaves (Newberry et al. n.d.) |
| *Xanthomonas perforans* AL65::Tn7-MRE152 | Xp; green fluorescent, chloramphenicol, kanamycin, and streptomycin resistant | This study |
| *Xanthomonas arboricola* CFBP 6826 | Xa WT | Pena et al. unpublished, tomato leaves |
| *Xanthomonas arboricola* CFBP 6826::Tn7-MRE160 | Xa; blue fluorescent, chloramphenicol, and tetracycline resistant | This study |
| *Pseudomonas capsici* 93B.260 | Pc WT | Laboratory collection, tomato leaves |
| *Pseudomonas capsici* 93B.260::Tn7-MRE145 | Pc; red fluorescent, chloramphenicol, and gentamicin resistant | This study |
| Plasmid | Features | Source |
| pMRE-Tn7-152 | green fluorescent sGFP2, Amp<sup>R</sup>, Cam<sup>R</sup>, Kan<sup>R</sup> | (Schlechter et al. 2018) |
| pMRE-Tn7-160 | blue fluorescent protein mTagBFP2, Amp<sup>R</sup>, Cam<sup>R</sup>, Tet<sup>R</sup> | (Schlechter et al. 2018) |
| pMRE-Tn7-145 | red fluorescent protein, mScarlet-I, Amp<sup>R</sup>, Cam<sup>R</sup>, Gent<sup>R</sup> | (Schlechter et al. 2018) |

Amp<sup>R</sup>, Cam<sup>R</sup>, Tet<sup>R</sup>, Gent<sup>R</sup>, Kan<sup>R</sup>: <sup>R</sup> indicates antibiotic resistance respectively.

### Tracking In planta populations of individual strains from mixed inoculation and disease severity measurement under high and low humidity

Bacterial cultures of overnight grown *Xanthomonas perforans* AL65::Tn7-mre152 (*Xp*, Km^R^), *Xanthomonas arboricola* CFBP 6826::Tn7-mre160 (*Xa*, Tet^R^) and *Pseudomonas* species 93B.260::Tn7-mre145 (*Pc*, Gm^R^) were suspended in MgSO_4_ buffer and adjusted to OD_600_ = 0.3, i.e., to a concentration of 10^8^ CFU/mL, and next the suspensions were diluted 100 times to a final concentration of 10^6^ CFU/mL. In this population study, 4–5-weeks-old tomato cv. FL 47R plants were inoculated by invertedly dipping for 30 seconds in 600 ml cell suspensions containing ∼1 × 10^6^ CFU/ml of *Xp* or *Xa* or *Pc* alone or *Xp* + *Pc* or *Xp* +*Xa* or *Xp* + *Xa* + *Pc* mixed in 1: 1 or 1: 1: 1 ratio for the mixed inoculation study. To understand the influence of humidity, each treatment was incubated under both low and high humidity. Each inoculum suspension was amended with 0.025% (vol/vol) of Silwet 77. Control plants were inoculated with sterile MgSO_4_ ammended with 0.025% (vol/vol) of Silwet 77. To understand the influence of humidity on bacterial population, each treatment combination was incubated under both low and high humidity. After dip inoculation, plants were kept inside closed plastic boxes, which were placed inside growth chambers with 12h light/dark cycle and 25°C temperature. Wet paper towels were placed at the bottom part of the walls of boxes for high humidity. The closed boxes containing the plants were not disturbed for the next 48 hours to maintain high humidity. After the first two days, lids were removed from the boxes containing plant replicates for low humidity conditions. For plant replicates with high humidity conditions, each day lids were taken off from the boxes for a 12-hour light cycle, and the boxes were closed back for the 12-hour dark cycle. Paper towels inside the boxes of high humidity conditions were continuously kept wet to maintain the humidity conditions. Leaf samples were collected after two hours, 3, 6, 9, and 14 days post-inoculation (dpi) from the plants. A sterile cork borer of radius ∼ 0.5 cm was used to collect 4 circular leaf discs from each leaf. Therefore, at each sampling point, ∼ 3 cm^2^ area of leaflet tissue was collected from each plant. A sterile forceps was used to place the leaf discs inside sterile microcentrifuge tubes containing 1 ml sterile MgSO_4_ buffer. The leaf tissue was macerated using a sterile homogenizer (Dremel). Serial dilutions of the homogenized leaf tissue suspension were plated on selective media using spiral plater (Neu-tec Group Inc, NY, USA) to estimate the population size of *Xp, Xa*, and *Pc* in individual and mixed inoculations. Plates were incubated for 3-4 days at 28°C before enumerating the number of colonies and bacterial populations was calculated as colony forming units (CFU) per cm^2^ of leaf area. Disease severity of bacterial leaf spot in the plants was measured using the following scale, and mean disease severity was calculated. Disease scale: 1 = symptomless, 2 = a few necrotic spots on a few leaflets, 3 = a few necrotic spots on many leaflets, 4 = many spots with coalescence on few leaflets, 5 = many spots with coalescence on many leaflets, 6 = severe disease and leaf defoliation, and 7= plant dead (Abbasi et al. 2002). Three plant replicates were used for each treatment. The experiment to track the in-planta population of *Xp* was repeated three times while In-planta population of *Xa* and *Pc* was enumerated in two of the three experimental sets.

### Biolog assay for the determination of nutritional similarity (NOI i.e. Nutritional Overlap Index)

Pure cultures of the bacterial strains were grown on BUG (biology universal growth) agar for 24 hours at 28°C. Three different microplates were used for three bacterial strains. The inoculum was suspended in inoculating fluid (IF-A). Cell density was adjusted to 95% transmittance. Next, the Biolog microplate (GEN III Microplate) was filled with 100 μl of the inoculum in each well. The plates were incubated for 48 hours at 28°C. Result was interpreted manually by observing the color change and measuring the optical density. Nutritional similarity was calculated with the formula NOIc = the number of carbon sources used by both the nonpathogenic bacterium and the pathogen/the total number of carbon sources used by the pathogen (Wilson and Lindow 1994b). This procedure was repeated thrice.

### Sequential infiltration assay to evaluate the influence of the interaction of *Pc* with the plant on symptom development and colonization by *Xp*

Five to six weeks old FL-47R tomato plants were placed in a growth chamber under ambient humidity, 12-hour light/dark cycle and day and night temperature of 28°C and 25°C respectively. For this experiment, bacterial strain *Xp* was used as effector-triggered susceptibility inducing “challenger” strain while *Pc* was used as PTI inducers and the experiment was followed similar to the PTI cell death assay described in Chakravarthy et al. (2009). All the bacterial strains were grown at 28°C for 48 hours followed by resuspending the cells separately in sterile MgSO_4_. The bacterial cells were washed twice followed by adjusting the final O.D of 0.3 (CFU 1 × 10^8^/ml) at 600 nm. The cell suspension was diluted 10 times using MgSO4 to get the final concentration of 1 × 10^7^ CFU /ml, which was used for infiltration. *Pc* was infiltrated on leaves in a rectangular area with a 1 ml sterile syringe. Five plants were inoculated with *Pc*. In each plant, 6 leaves were used for infiltration. 4 hours after the inducer infiltration, the leaves were infiltrated with *Xp*, in the same order of plants. A point on the periphery of the first inoculation rectangle was be used as the center of the second inoculation rectangle. The plants were placed back inside the growth chamber and monitored for the appearance of cell death in the area which were challenged with *Xp*. Cell death inside the overlapping region of infiltration indicated a breakdown of innate immune response by *Xp* (Chakravarthy et al. 2009).

### Data Analysis

Analysis of disease development using disease severity index, area under disease progress curve (AUDPC), and area under growth progress curve (AUGPC) was conducted and statistical significance testing was performed on linear mixed effects model. Treatments were used as fixed variable and experiments were used as random variable. Data analysis and visualization was done using mixed linear model in R following the scripts by (Schandry 2017).

## Results

### Mixed infection with *Xa* and *Pc* exacerbates disease severity on *Xp* infected tomato plants under high humidity conditions

The disease severity on tomato leaves co-inoculated with *Xp* +*Pc* +*Xa* displayed highest disease severity compared to double *Xp*+*Pc* or *Xp*+*Xa* or single infections (Figure 1).

**Figure 1.**
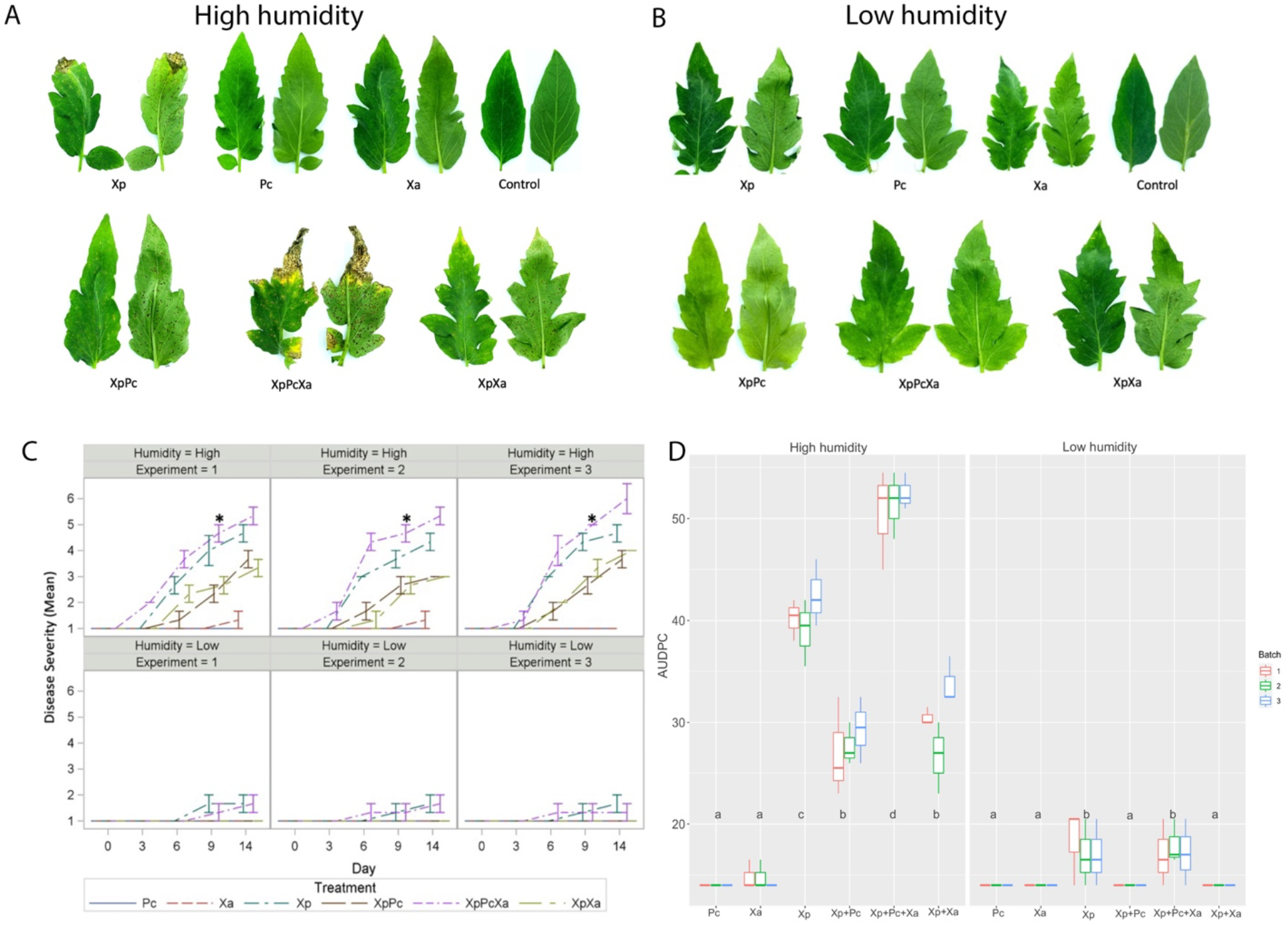
Disease Severity under low and high humidity in individual and mixed inoculated plants. Four to five weeks old tomato (cv. FL47) plants were inoculated with ∼1× 10^6^ cfu/ml of *Xp, Pc, Xa, Xp* + *Pc, Xp* + *Pc* + *Xa, Xp* + *Xa*. (a) Disease severity on day 14 under high humidity (b) Disease severity on day 14 under low humidity. (c) Disease severity over the 14-day duration of the experiment was plotted for individual inoculation and mixed inoculation treatments from three different experimental batches. Vertical lines represent the 95% confidence limit. ANOVA repeated measurements (GLIMMIX model) wereere applied to analyze the disease severity values statistically. Treatment *XpPcXa* with * mark on day 9 is significantly higher disease severity compared to *Pc, Xa, Xp* + *Pc, Xp* + *Pc* + *Xa, Xp* + *Xa* according to Tukey’s test of least significant difference (P<0.05) (d) Area under disease progress curve (AUDPC) for tomato plants co-inoculated with the different treatments is plotted. AUDPC raw values and mean with grouping letters (a, b, c, and d) according to significant difference (*p* value<0.05) from the linear mixed model with 95% confidence interval per treatment plotted from three different experimental batches.

Humidity influenced the disease severity levels associated with both the individual and mixed infections, with high humidity supporting higher disease severity levels in single or mixed inoculations. Bacterial spot symptoms, recorded as water-soaking, necrotic and/ chlorotic lesions, appeared as early as 4 dpi under high humidity compared to 6 dpi under low humidity (Figure 1A, B). Plants inoculated with *Xa*, or *Pc* alone did not show any significant differences in disease severity between high and low humidity. *Pc* did not develop any visible symptoms under either high or low humidity condition throughout the 14 days of experiment (Figure 1A, B). Plants inoculated with *Xa* developed some symptoms only under high humidity after 9 dpi. On 9 dpi, under high humidity, plants co-inoculated with the three bacterial species *Xp, Pc*, and *Xa* showed significantly higher mean disease severity (p<0.05) ranging from 5 to 6 on a disease scale, compared to *Xp* + *Pc* and *Xp + Xa* or *Pc* and *Xa* alone showing disease severity between 1 to 2.5 on the disease scale (Figure 1C). Under low humidity, there were no significant differences in disease severity among any of the treatments over the 14-day period. We quantified the differential virulence in single vs mixed infection by comparing the area under the disease progress curve (AUDPC) for each treatment under low and high humidity conditions. The highest AUDPC values (Figure 1D) were observed in case of mixed infection of *Xp, Pc* and *Xa*, followed by *Xp* alone under high humidity conditions (p <0.01). *Xp* when co-inoculated with *Pc* or *Xa* showed lower AUDPC compared to *Xp* alone under high humidity conditions. Under low humidity, *Xp* and *Xp* + *Pc* + *Xa* showed significantly higher disease severity compared to other treatments (p <0.01). Together, these results demonstrate that presence of weak or opportunistic pathogens can alter *Xp* disease dynamics in a humidity-dependent manner.

### *Xp* population is predominantly reduced in mixed infection treatments

To understand the influence of co-infection on *Xp* in planta growth dynamics, we next measured *Xp* population over time in different treatments. Presence of *Pc* lowered *Xp* population significantly compared to individual *Xp* infection under both low and high humidity conditions during later stages of the disease development, i.e. 9 dpi (p_high_ <0.05; p_low_ < 0.01) and 14 dpi (p_high_=0.01, p_low_ < 0.05) (Figure 2). Similar observations of the impact of mixed infection of *Xp +Xa+Pc* on *Xp* population were noted under high humidity (p=0.05 at 9dpi and p <0.001 at 14 dpi), but not under low humidity. In contrast to *Pc* having influence on later stage population dynamics, presence of *Xa* significantly lowered the population of *Xp* more than one log compared to *Xp* alone throughout the course of the experiment (Figure 2A) under high humidity conditions (p < 0.001). We confirmed these observations by comparing the area under growth progress curves (AUGPC) that indicated the significant influence of co-infection on *Xp* growth dynamics under both high and low humidity (Figure 2B). However, co-infection with *Xa* appeared to drastically reduce *Xp* growth by lowest AUGPC values compared to *Xp* + *Pc*+ *Xa* or *Xp* +*Pc* treatments. Together, these results suggest that co-infection reduces overall *Xp* population despite higher disease severity and that different co-infecting species may differentially influence the growth dynamics of *Xp*.

**Figure 2.**
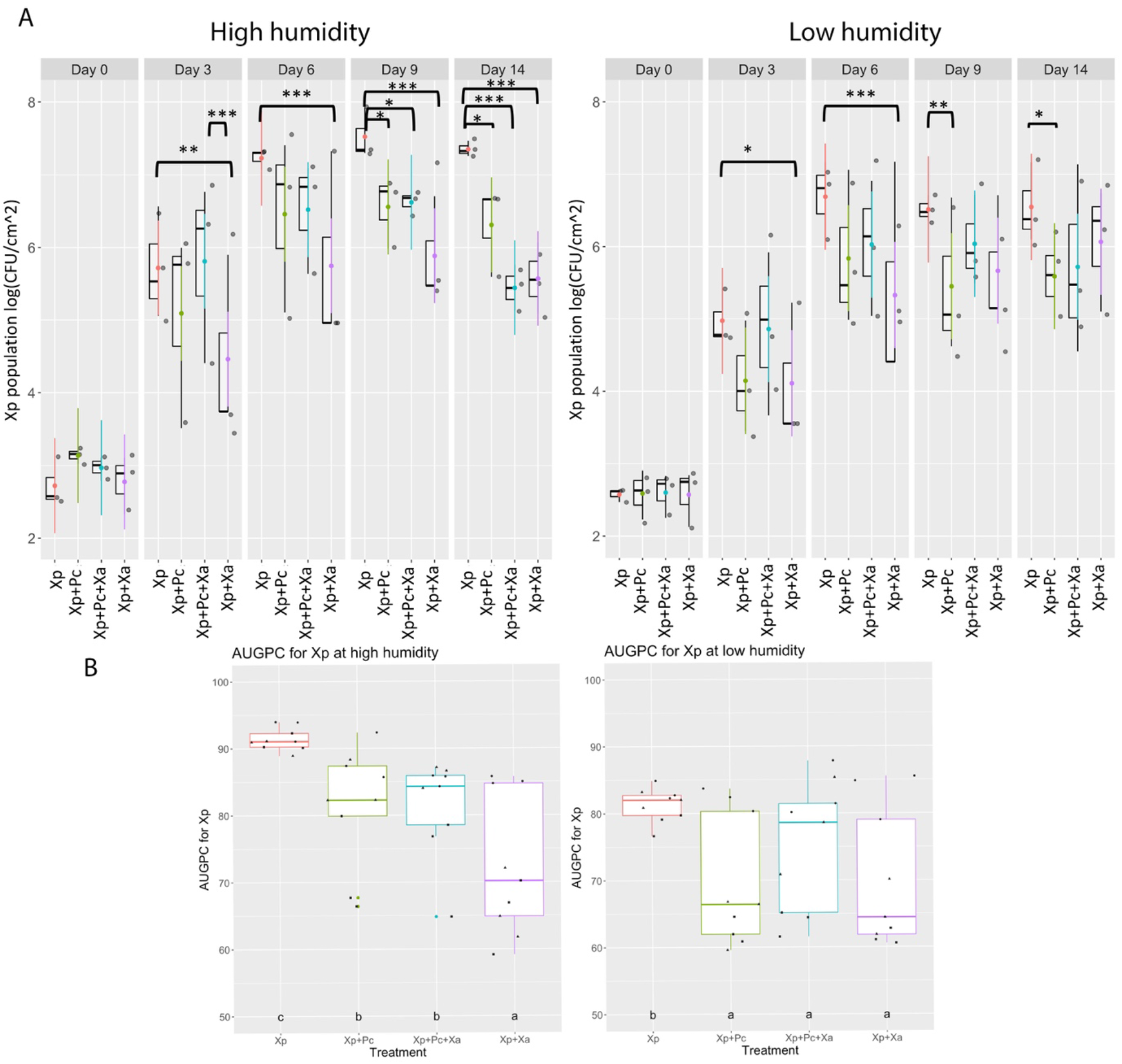
Effect of mixed inoculation on *in planta* population dynamics of *Xp*. Four to five weeks old tomato (cv. FL47) plants were inoculated with ∼1× 10^6^ cfu/ml of *Xp, Xp + Pc, Xp + Xa*, and *Xp + Xa + Pc* and maintained under high and low humidity conditions. The experiment was repeated three times and experiment was considered as a batch factor in linear mixed model. (A) Growth of *Xp* population was evaluated from plants inoculated with the different treatments on day 0, day 3, day 6, day 9 and day 14 post inoculation on selective media. (A) ANOVA (mixed linear model) was applied for the statistical analysis of the log_10_ cfu/cm^2^ of *Xp* values. Significant differences (P<0.05) among the treatments, according to Tukey’s test of least significant difference are depicted in the graph by asterisks. * represents p<0.05; ** as p<0.005; *** as p<0.0005. (B) Area under growth progress curve (AUGPC) raw values and mean with grouping letters (a, b, c and d) according to significant difference (*p* value<0.05) from linear mixed model with 95% confidence interval per treatments plotted from three different experimental batches. Each experimental batch is indicated by a different shape.

### A significant increase in *Xa* population is observed under co-infection

We observed small lesions on *Xa* infected plants under high humidity, similar to previous reports of *Xa* being opportunistic or weak pathogen on tomato under field conditions (Mbega et al. 2012a). Here, we assessed whether *Xa* is truly opportunistic pathogen when coinfected with *Xp* or *Xp* +*Pc* and whether humidity had influence on the opportunistic nature of *Xa*. High humidity did not significantly alter AUGPC values for Xa compared to low humidity when Xa was present alone on the leaves, although overall Xa population was ∼ 10-50-fold higher under high humidity compared to low humidity (Figure 3A). Variability in AUGPC values as well as population levels each day was observed at low humidity when Xa was present alone, which could indicate that Xa population is highly susceptible to environmental conditions. However, co-infection with *Xp* significantly promoted Xa growth throughout the experiment under both high and low humidity conditions (p < 0.001), except for 14dpi at low humidity when *Xa* populations were lowest in all treatments. Similarly, *Xp+Pc+Xa* co-infection led to higher *Xa* population with higher AUGPC compared to *Xa* alone under both high and low humidity conditions (Figure 3B). This three-species mixed infection resulted in similar AUGPC values to *Xa+Xp* under high humidity, but slightly lower compared to *Xa* +*Xp* under low humidity. Presence of *Pc* altered *Xa* population when in co-occurrence with *Xp* under low humidity conditions.

**Figure 3.**
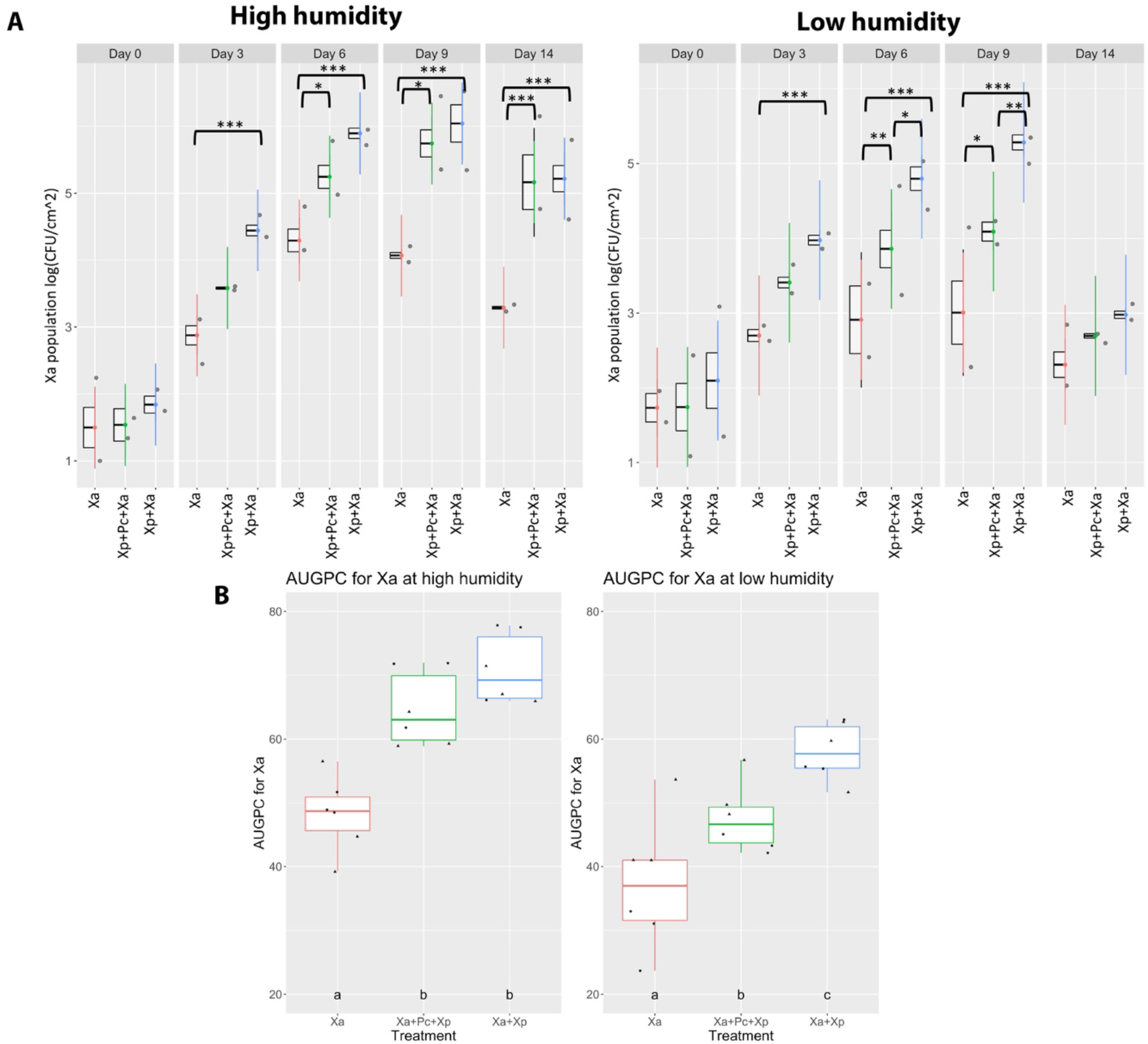
Effect of *Xp* and *Pc* on *Xa in planta* population. Four to five weeks old tomato (cv. FL47) plants were inoculated with ∼1× 10^6^ cfu/ml of *Xa, Xp + Pc* + *Xa, Xp + Xa* and incubated in growth chamber under high and low humidity conditions. The experiment was repeated three times and experiment was considered as a batch factor in linear mixed model. (A) Growth of *Xa* population was evaluated from plants inoculated with the different treatments on day 0, day 3, day 6, day 9 and day 14 post inoculation on selective media. ANOVA (mixed linear model) was applied for the statistical analysis of the log_10_ cfu/cm^2^ of *Xp* values. Significant differences (*p*<0.05) among the treatments, according to Tukey’s test of least significant difference are depicted in the graph. (B) Area under growth progress curve (AUGPC) raw values and mean with grouping letters (a, b, c and d) according to significant difference (*p* value<0.05) from linear mixed model with 95% confidence interval per treatments plotted from three different experimental batches. Each experimental batch is indicated by a different shape.

### *Pc* population is higher in presence of mixed infection with Xp and Xa

No visual disease symptoms were observed when *Pc* was inoculated alone on tomato leaves. *Pc* by itself grows poorly on tomato leaves, although high humidity can support 10-50-fold higher growth compared to low humidity conditions. In presence of *Xp* and *Xp +Xa, Pc* population was 10-100-fold higher compared to *Pc* alone, both under high and low humidity conditions (p <0.01) (Figure 4A and B). The influence of humidity was observed when comparing three-species infection and two-species infection, where absence of *Xa* led to higher AUGPC values of *Pc* population under high humidity.

**Figure 4:**
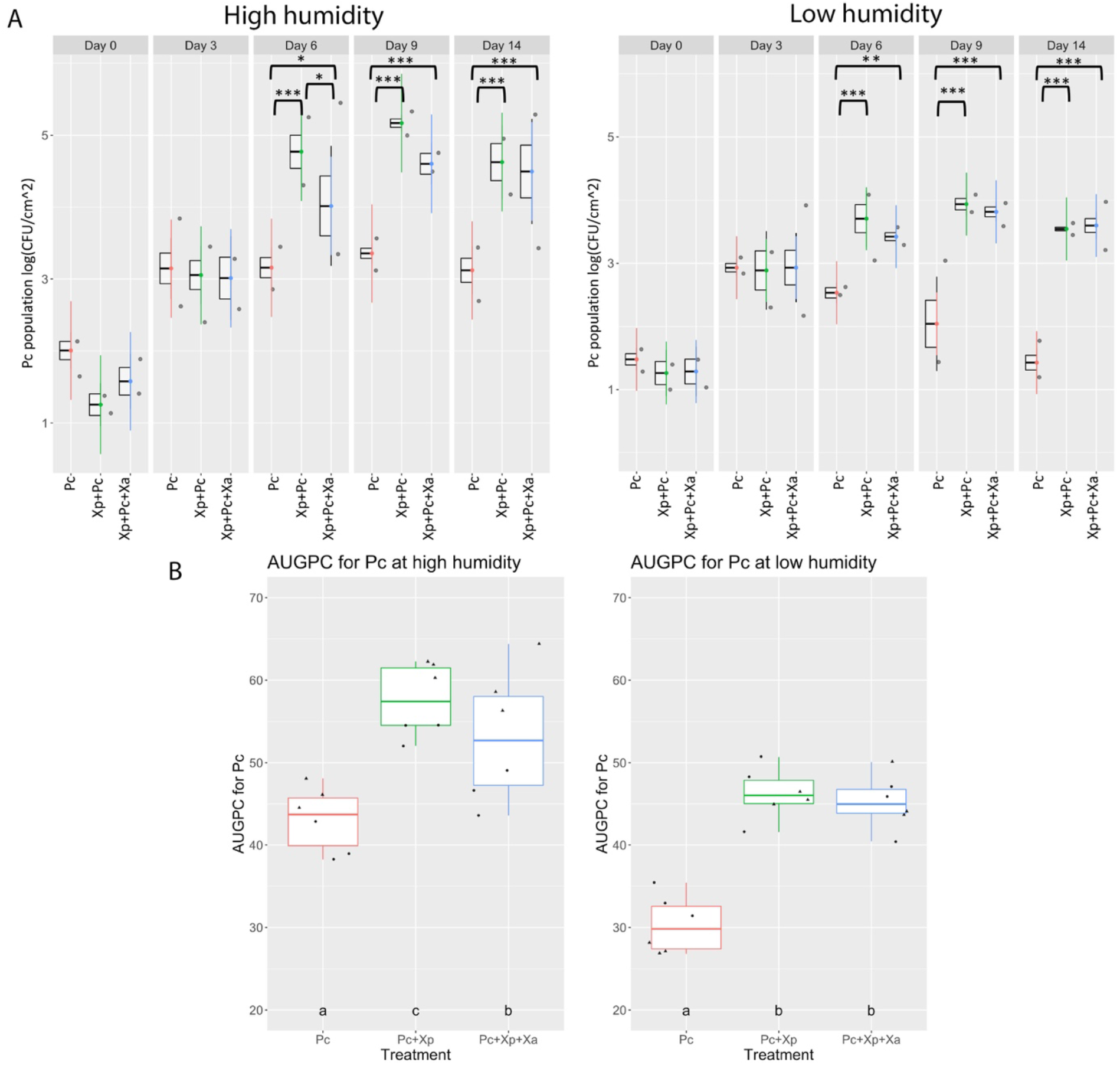
Influence of *Xp* and *Xa* on *Pc in planta* population. Four to five weeks old tomato (cv. FL47) plants were inoculated with ∼1× 10^6^ cfu/ml of *Pc, Xp + Pc, Xp + Pc + Xa* and incubated in growth chamber under high and low humidity conditions. The experiment was repeated three times and experiment was considered as a batch factor in linear mixed model. (A) Growth of *Pc* population was evaluated from plants inoculated with the different treatments on day 0, day 3, day 6, day 9 and day 14 post inoculation on selective media. ANOVA (mixed linear model) was applied for the statistical analysis of the log_10_ cfu/cm^2^ of *Xp* values. Significant differences (*p* < 0.05) among the treatments, according to Tukey’s test of least significant difference are depicted in the graph. (B) Area under growth progress curve (AUGPC) raw values and mean with grouping letters (a, b, c and d) according to significant difference (*p* value<0.05) from linear mixed model with 95% confidence interval per treatments plotted from three different experimental batches. Each experimental batch is indicated by a different shape.

### Higher nutritional similarity (NOI_C_) between *Xp* and *Xa*

To understand if the altered *Xp* population and AUGPC in presence of the co-occurring bacterial species was an outcome of nutritional resource competition, we determined nutritional overlapping index (NOI) i.e. nutritional similarity among three strains using the carbon source utilization profiles (Supplementary figure 1), according to Wilson and Lindow (Wilson and Lindow 1994a). NOIc = the number of carbon sources used by both the nonpathogenic bacterium and the pathogen/the total number of carbon sources used by the pathogen. The highest NOI value is 1.0, and it indicates higher nutritional similarity between the two bacterial species. The carbon utilization profile between *Xp & Xa* is 0.96, which is higher than the others. Niche overlap values greater than 0·9 is indicative of utilization of the same nutrients (Lee and Magan 1999; Wilson and Lindow 1994b). The NOIc between *Xa & Pc* is 0.84 and the NOIc between *Xp & Pc* is 0.81. The results indicated that there is more nutritional overlap among *Xa* and *Xp* but lower among *Pc* and *Xp*.

### Pre-infiltration of tomato leaves with *Pc* delays cell death caused by *Xp*

Since we ruled out nutritional similarity being a reason for lower *Xp* population in presence of *Pc*, we conducted sequential inoculation of *Pc* and *Xp* using infiltration directly into apoplast to test the hypothesis that priming of plants by *Pc* leads to delay in symptom development and growth suppression of *Xp* in planta. *Pc* by itself grows poorly in planta and causes minimal or no obvious symptoms when dip-inoculated or directly infiltrated into the apoplast. *Xp* was infiltrated 4 hours after infiltration of *Pc* such that symptom development by *Xp* and growth of *Xp* can be tracked in the *Xp* alone and sequential *Pc*-*Xp* infiltration area. Water-soaking phenotype was observed in the area infiltrated with only *Xp* (Figure 5a). There were no symptoms in the overlapping region of the leaf infiltrated with both *Pc* and *Xp* or *Pc* alone. After the next 24 hours i.e. day 4 after inoculation, the symptoms developed in the overlapping region. Thus, a delay in cell death symptoms by *Xp* was observed in the overlapping area where prior *Pc* infiltration was carried out 4 hours before challenge inoculation with *Xp*, indicating delayed susceptibility response in presence of *Pc*. Next, we assessed whether delayed symptom development was also associated with lower *Xp* population in the overlapping region with sequential infiltration compared to *Xp* alone. Population of *Xp* from the overlapping region was half a log lower than *Xp* population from the region infiltrated with only *Xp* (Figure 5b). However, the difference in the population was not significant. Infiltration with *Xa* in tomato apoplast led to cell-death reaction, which prevented us from evaluating influence of pre-infiltration of *Xa* on *Xp* symptom development (Supplementary Figure 2). Collectively, these results indicate that *Pc* may elicit defense response in the form of innate immunity (as observed as early as 4 hours upon infiltration) that led to delayed symptom development by *Xp*.

**Figure 5:**
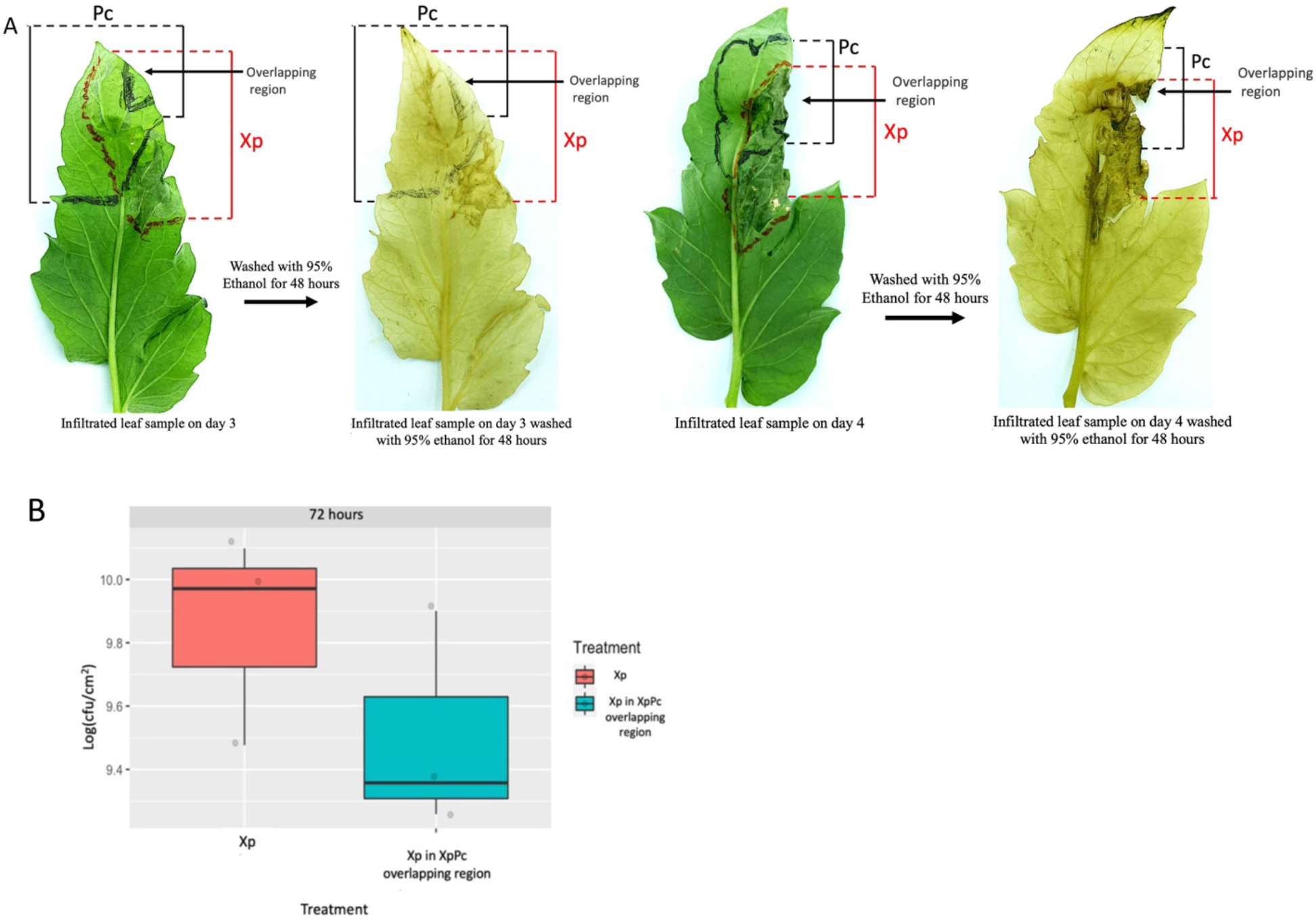
Sequential infiltration assay to assess influence of plant innate immunity on bacterial spot symptom development and *Xp* population. Tomato leaves were infiltrated with *Pc* (area between the two black lines), and 4 hours later, the leaves were challenged with *Xp* (area between the two red lines). (A) On day 3, water soaking or cell death symptoms by *Xp* were observed in the *Xp* alone infiltrated region. No cell death was observed in the overlapping regions. Cell death was clearly visible after ethanol wash. On day 4, cell death was observed in the overlapping area. (B) Day 3 population of *Xp* from the overlapping region infiltrated with *Xp + Pc*, and population of *Xp* from the area infiltrated with *Xp* only. Vertical lines represent standard error bars. t-test was done on log10 cfu/cm^2^ values and difference between two treatments were not significant (P<0.05). Concentration of each inoculum was 1 × 10^7^ CFU /ml.

## Discussion

This study provides empirical evidence that co-existing weak pathogens can influence within-host disease dynamics and that the humidity can play an important role in co-infection dynamics. We specifically used foliar bacterial pathogen, *Xanthomonas perforans*, endemic to many tomato-growing regions. The continued disease outbreaks are common for this pathogen. Different biotic and abiotic factors are thought to drive the severity of the disease outbreaks. However, systematic assessment of these factors using empirical approaches is lacking. Our co-inoculation and sequential inoculation experiments confirm that co-infection by multiple species can change the disease load (Figure 1) and interspecies interaction can influence disease and Xp growth dynamics

(Figure 1, 2). Humidity was found to be a significant factor in determining the overall fate of the interspecies interactions (Figure 2, 3, 4). These results confirm that high humidity can favor higher disease severity, not by supporting higher *Xanthomonas perforans* population, but rather by also fostering growth of opportunistic pathogens or weak pathogens.

It has been observed that *Xanthomonas perforans* requires high humidity for enhanced disease development and tropical or subtropical regions with high humidity has been previously identified to have the ideal condition for them (Abrahamian et al. 2021; Carvalho et al. 2019; Obradovic et al. 2008). Results of our study substantiate this observation based on higher disease severity when compared under high and low humidity conditions. Co-infection with *Xp, Pc* and *Xa* favored higher disease severity compared to *Xp* alone. Tracking within-host population revealed that higher disease severity under co-infection is not due to higher *Xp* population. Rather, we observed that co-infection with all three species favored higher growth of *Pc* and *Xa* under high humidity conditions. High humidity did not significantly induce symptom development in plants inoculated with either only *Pc* or only *Xa*, although we observed spot-like symptoms on *Xa* inoculated plants at the late stages of time course experiment. Different pathovars of *Xanthomonas arboricola* have been observed to cause disease in humid and warm condition (Kałużna et al. 2021; Lamichhane 2014; Lamichhane and Varvaro 2014). This observation suggests opportunistic pathogenic nature of *Xa* during its interaction with tomato plants. Similar benefit for higher *Pc* survival in the phyllosphere was observed based on in planta population similar to previous studies (Cambra et al. 2004; Janse 1987). *Pseudomonas cichorii*-like strains has been recorded to cause pith necrosis in tomato plants (Trantas et al. 2013). Several commensal *Pseudomonas* spp. like *P. marginalis, P. putida, P. protegens, P. citronellolis* have been observed to cause increased disease severity of pith necrosis in tomato when coinfected with *Xp* (Aiello et al. 2017). However, in our study when we examined for pith necrosis symptoms by *Pc*, which is closely related to *P. cichorii*, we did not observe any vascular browning in the plants that were inoculated with either *Pc* or both *Xp & Pc* (data not shown).

Interestingly, influence of co-infection also seemed to depend on the coinfecting member, regardless of the humidity conditions. Co-infection by *Xa* or *Pc* with *Xp* led to lower AUDPC values compared to *Xp* alone or co-infection by all three coinfecting members. To understand the differential impact of coinfecting members on disease severity, we further compared population dynamics of individual coinfecting members. As expected, highest AUGPC value for *Xp* population was observed when *Xp* was alone, followed by *Xp +Pc* or *Xp +Pc +Xa* followed by lowest value when *Xp* was coinfected with *Xa*. Indeed, *Xa* benefits from co-infection with *Xp* as indicated by *in planta* population and AUGPC values. We further compared nutritional overlapping index to assess resource competition experienced during co-infection. This experiment indicated that *Xp* and *Xa* likely compete for resources which results in lower *Xp* population when in presence of *Xa. X. arboricola* has been indicated as opportunistic pathogen and in some cases, putative causal agent in bacterial spot infected tomato and pepper fields (Roach et al. 2018). *Xa* strain CFBP 6826 used in this study contains type III secretion system as well as type III effectors, which may induce susceptibility reaction when gained access to apoplastic space in presence of *Xp* (Pena et al. Unpublished). Further experiments to study colonization success of *Xa* in mixed infection and alone may explain whether *Xa* has ability to gain access to apoplastic space by itself. *In planta* population of *Xa* when inoculated by itself did not lead to significant increase in its population. Nutritional overlap did not explain lower *Xp* population in presence of Pc. Rather, we observed that pre-treatment with *Pc* resulted in delayed symptom development by *Xp* and reduced *Xp* population. This may suggest that interaction of *Pc* with the plant results in possible induction of defense response. *Pc* by itself may not be able to overcome the innate immune response, thus, explaining why *Pc* population remains low when inoculated alone. However, suppression of plant innate immune response by *Xp* may allow *Pc* to grow to higher populations. However, co-infection with *Pc* lowers *Xp* populations. Assuming *Xa* competes with *Xp* for resources in dual co-infection treatment and lowers *Xp* populations, our observation of higher *Xp* population in three-species co-infection treatment compared to dual-species co-infection with *Xa* was surprising. The influence of resource competition is minimal when *Xp* colonizes along with *Pc* and *Xa*, as indicated by higher *Xp* population in presence of all three coinfecting species compared to *Xp +Xa*. This may further suggest potential synergism among *Xp* and weak pathogen, *Xa*, under high humidity conditions, where behavior and expression of virulence factors such as type III effectors, or type II secreted cell-wall degrading enzymes might be synchronized. Such synchronization mediated by sharing of quorum sensing signals among *Pseudomonas savastanoi* and nonpathogenic species, *E. toletana* and *P. agglomerans* was shown in olive-knot disease, a classic example of poly-bacterial disease (Hosni et al. 2011). We speculate that the induction of innate immune response by *Pc* may reduce resource competition among *Xp* and *Xa* and rather enhance synergistic interactions among *Xp* and *Xa*, thus, allowing *Xp* to maintain higher population in presence of all three coinfecting species compared to *Xp +Xa*.

Overall, we can conclude that dominant bacterial leaf spot pathogen *Xanthomonas perforans* population depends on the presence of its co-occurring bacterial species *Pc* and *Xa*, but also on environmental conditions, specifically, humidity. We observed that humidity can alter interspecies interactions. Humidity gives advantage to these two weak pathogens to colonize better and reduce population of *Xp*. These three bacterial species compete for resources and spatial distribution in the leaf phyllosphere which in turn results in lower carrying capacity and lower population of *Xp*. The plant’s immune response raised by *Pc* also managed to delay the cell death susceptibility of *Xp* and caused delayed symptom development. In fields, when the weather is warm and humid, we can expect to see higher bacterial spot disease severity in presence of the co-occurring bacterial species compared to if the dominant pathogen was present alone. *Xp* makes it easier for *Xa* and *Pc* to colonize the host plant, but their presence becomes detrimental for the *Xp* population and reduces it. This study highlights the importance of community interactions and co-occurring weak pathogens on overall disease outcome, the unexplored area in plant disease studies. Co-infection can be an important driver of epidemiological dynamics, leading to increased transmission (Susi et al. 2015) and thus, more widespread disease outbreaks. Thus, further studies investigating epidemiological impact, specifically on transmission of principal pathogen, *Xp*, may help explain impact of co-occurring weak pathogens on field level pathogen dynamics. This is also important consideration for disease control programs that may need to take into account co-occurring weak pathogens.

## Acknowledgements

We thank Plant Science Research Center staff at Auburn University for providing greenhouse space and maintenance of plants used for experiments in this study. We also thank CIRM-CFBP (Beaucouzé, INRAE, France https://cirm-cfbp.fr/) for Xa strain preservation and supply.

---

**Supplementary Figure 1.**
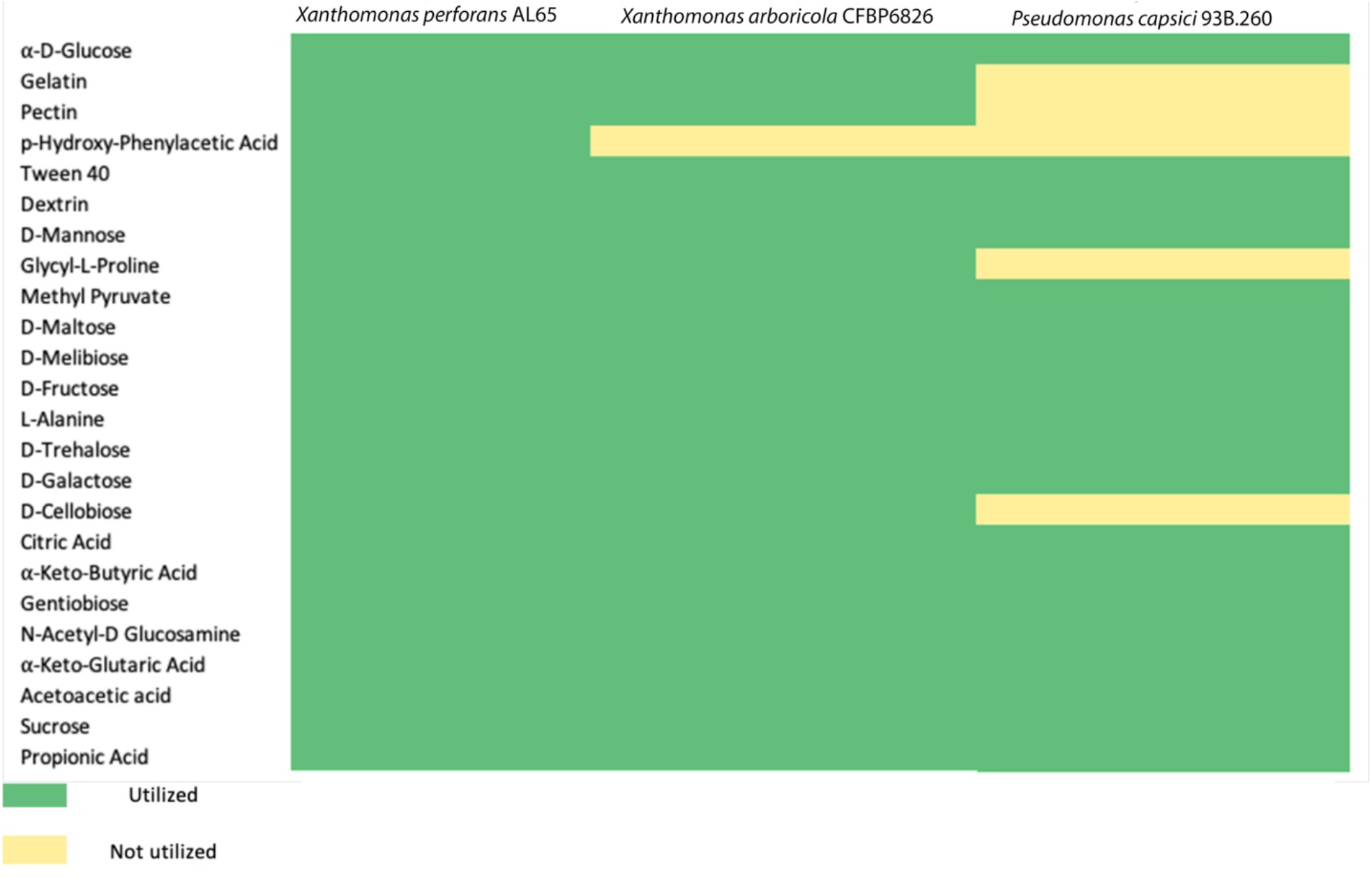
Carbon source utilization Profile of *Xanthomonas perforans* AL65, *Xanthomonas arboricola* CFBP 6826, *Pseudomonas capsici* 93B.260

**Supplementary Figure 2.**
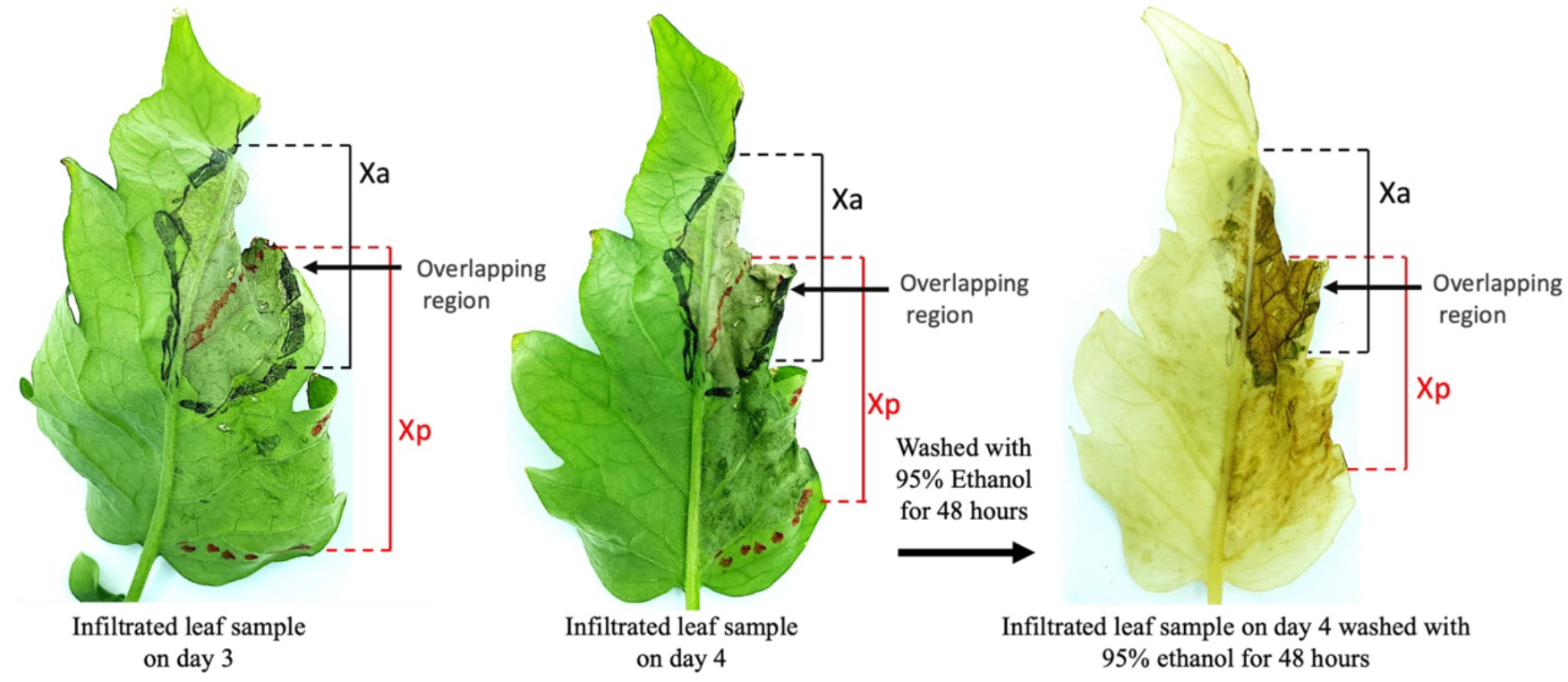
Tomato leaves were infiltrated with *Xa* to induce PAMP triggered immunity (area between the two black lines), and 4 hours later, the leaves were challenged with *Xp* (area between the two red lines). On day 3, water soaking or cell death symptom by *Xa* was observed in leaf area infiltrated with *Xa* and the overlapping region infiltrated with both *Xa* and *Xp*. On day 4, cell death was also observed in the area infiltrated with only *Xp*. Leaf with symptom from day 4 was ethanol washed to visualize the cell death better. Concentration of each inoculum was 1 × 10^7 CFU /ml.

